# CiliaMiner: an integrated database for Ciliopathy Genes and Ciliopathies

**DOI:** 10.1101/2022.11.28.518070

**Authors:** Merve Gül Turan, Mehmet Emin Orhan, Sebiha Cevik, Oktay I. Kaplan

## Abstract

Cilia are found in eukaryotic species ranging from single-celled organisms, such as *Chlamydomonas reinhardtii*, to humans, but not in plants. The ability to respond to repellents and/or attractants, regulate cell proliferation and differentiation, and provide cellular mobility are just a few examples of how crucial cilia are to cells and organisms. Over 30 distinct rare disorders generally known as ciliopathy are caused by abnormalities or functional impairments in cilia and cilia-related compartments. Because of the complexity of ciliopathies and the rising number of ciliopathies and ciliopathy genes, a ciliopathy-oriented and up-to-date database is required. In addition, disorders not yet known as ciliopathy but have genes that produce cilia localizing proteins have yet to be classified. Here we present CiliaMiner, a manually curated ciliopathy database that includes ciliopathy lists collected from articles and databases. Analysis reveals that there are 55 distinct disorders likely related to ciliopathy, with over 4000 clinical manifestations. Based on comparative symptom analysis and subcellular localization data, diseases are classified as primary, secondary, or atypical ciliopathies. CiliaMiner provides easy access to all of these diseases and disease genes, as well as clinical features and gene-specific clinical features, as well as subcellular localization of each protein. Additionally, the orthologs of disease genes are also provided for mice, zebrafish, Xenopus, Drosophila, and *C. elegans*. CiliaMiner (https://kaplanlab.shinyapps.io/ciliaminer) aims to serve the cilia community with its comprehensive content, and highly enriched interactive heatmaps, and will be continually updated.

## Introduction

Cilia are cellular organelles that extend out of the cell in unicellular and multi-organismal organisms. Within the cilia structure, nine pairs of peripheral microtubules are radially positioned and encircled by the cilia membrane, and this microtubule-based core structure is called the axoneme. Despite cilia being tiny cellular organelles, cilia have several subcompartments, including the transition zone (TZ), the basal body (a modified centriole), the axoneme, and the distal segment (1). Importantly, the ciliary structures and subcompartments have been well-preserved throughout evolution. Furthermore, depending on their structural differences and functional distinctions, cilia are categorized as either motile (9+2 axonemal structures) or non-motile (9+0 axonemal structure; primary cilium) **(Figure 1)** (2). In humans, motile cilia exist as multiple cilia on cell surfaces, requiring cell motility and fluid movement, whereas a single non-motile cilium emerges from a cell surface and is primarily involved in sensation (chemosensation and photosensation), developmental and signaling pathway regulation (Wnt, Hedgehog) (3,4). Any structural or functional anomalies in primary or motile cilia can cause a variety of rare disorders, and the term “ciliopathy” can be used to describe the complete spectrum of diseases that are related to primary or motile cilia (5).

**Figure 1:**
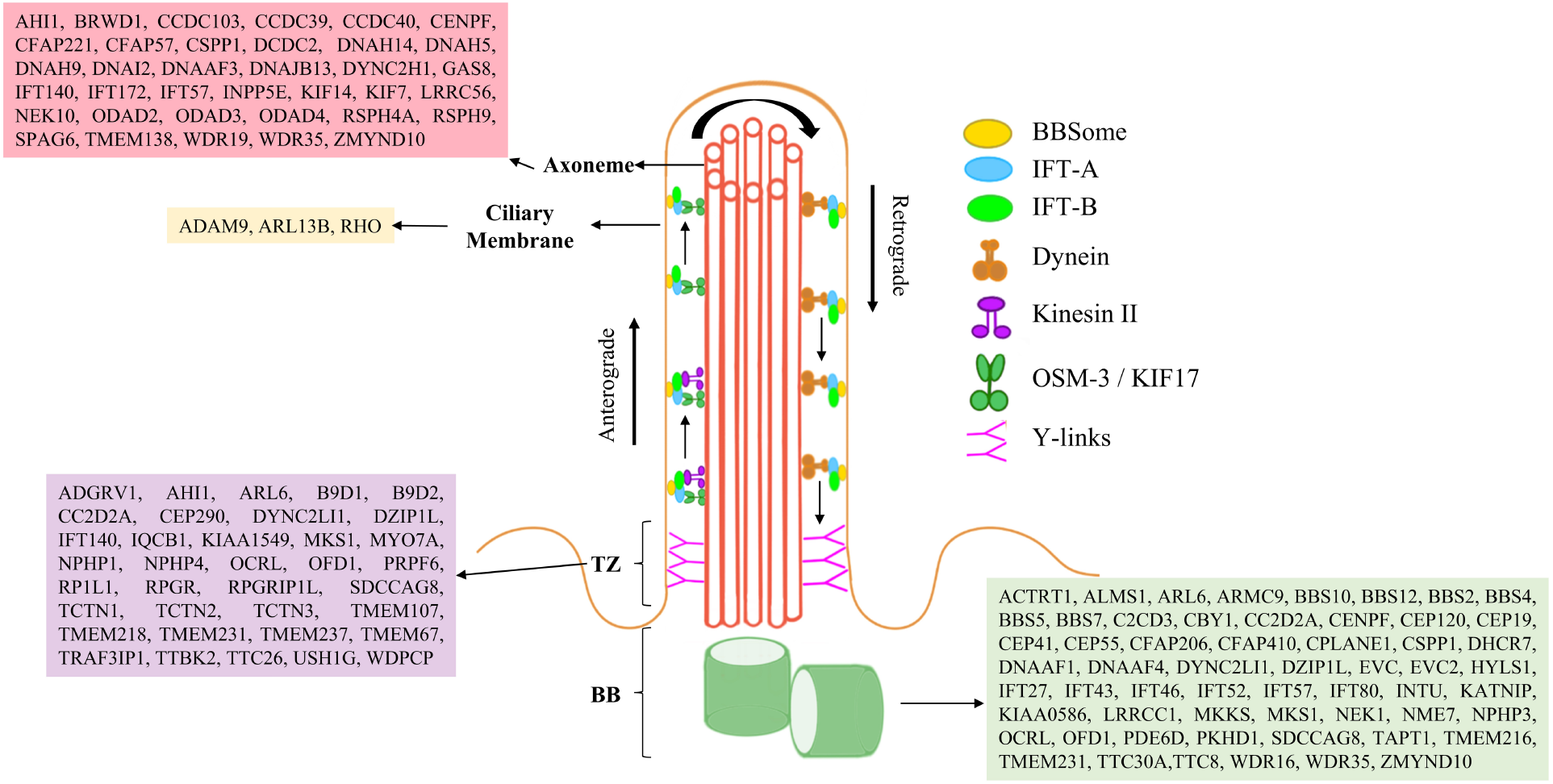
The primary cilium is depicted, with a 9 + 0 microtubular axoneme surrounded by a ciliary membrane. Two ciliary subcompartments, including the basal body (BB) and transition zone (Y-shaped linkers) are shown. Intraflagellar transport (IFT), including motor proteins (kinesin and dynein), IFT-A, IFT-B, and Bardet-Biedl syndrome proteins (BBSome) move in both directions (anterograde and retrograde) along cilia. Ciliary localization of proteins encoded by primary and secondary ciliopathy-causing genes is shown in a representative cilia structure.

The number of ciliopathies and ciliopathy-related genes has expanded dramatically over the last 20 years, owing to advances in medical technology, the ease of disseminating clinical knowledge, and the ability to sequence people (6). Because of the growing number of ciliopathies and disease-associated genes, compiling, categorizing, and displaying the numerous ciliopathies and their associated genes has become more difficult. Furthermore, symptomatic differences between ciliopathies, changes in symptom frequency, and the same gene causing multiple types of ciliopathies complicate the issue. Even though all the relevant details and information about diseases and disease genes are available, they are scattered among different biological and medical databases, necessitating users to visit these databases like MalaCards (7) and OMIM (8). It is critical to keep all of this massive data up to date and in one place, allowing the ciliopathy community to access all of the most recent data.

Here we present a ciliopathy-specific database, called CiliaMiner, that compiles and presents the updated list of ciliopathies, ciliopathy-associated genes, and the disease symptoms for each condition. Furthermore, CiliaMiner provides classifications of ciliopathies and associated disorders based on the subcellular localization and functions of disease-associated genes in conjunction with disease symptoms. CiliaMiner offers easy access to all ciliopathies, disease symptoms, ciliopathy genes, ciliopathy candidate genes, and orthologs of ciliopathy genes and ciliopathy candidate genes, seeking to serve as a major place in the ciliopathy database. Each piece of information is manually acquired and confirmed before being uploaded to the appropriate part of the CiliaMiner database, and users can also easily submit their data to CiliaMiner.

## Materials and Methods

### Data collection and curation

Data presented in the CiliaMiner were manually collected from six separate data sources, including Online Mendelian Inheritance in Man (OMIM) (8), PubMed, Congruent clinical Variation Visualization Tool (ConVarT) (9), OrthoList 2 (OL2) (10), Wormbase (11), and Protein Atlas (12) databases. We specifically utilized the phrase “ciliopathy” in PubMed to find a comprehensive list of ciliopathy disorders, followed by confirmation of ciliopathy disease using an OMIM search. The manual search, together with OMIM validation, yielded 55 possible ciliopathy diseases. Once the list of ciliopathy diseases is finalized, we collected the clinical symptoms and features of each disease using OMIM and PubMed web pages. The disease-associated genes and relevant references were collected from OMIM and PubMed. Additionally, we looked for potential ciliopathy genes in PubMed using the phrase “cilia” between 2018-2022, and if the protein product of the gene localizes to cilia, we next searched for disease relevance and collected symptoms for that disease. Finally, we visited PubMed for each gene to determine whether the protein encoded by a ciliopathy gene is detected in cilia or cilia-related compartments such as the basal body and transition zone. Finally, we obtained the localization from ProteinAtlas (12), an excellent source for protein localization, for those for which we were unable to find it.

### Orthologs of Disease-Associated Genes

The ciliopathy disease list was created using human genes. Following that, the ortholog genes of other organisms, *Mus musculus, Danio rerio, Xenopus laevis, Drosophila melanogaster*, and *Caenorhabditis elegans*, are created using ConVarT (9) and OrthoList 2 (10) (just for *C. elegans*). Wormbase used to be certain of the common gene name, sequence number, and WormBase ID of *C. elegans*. We created a dedicated webpage for orthology search, called “Ciliopathy Genes and Orthologs”.

### Collection of Detailed Clinical Symptoms of Diseases

Following the collection of the list of ciliary diseases, 4092 distinct clinical features are discovered for disorders connected to ciliopathy (OMIM and the articles). A meticulous collection of clinical symptoms suggests that there are 2354 and 1784 unique symptoms for primary ciliopathy and secondary disease groups, respectively. Because the list of clinical features is extensive, we choose the limited numbers of clinical symptoms to generate representative heatmaps on the “Ciliopathy Names” page in the CiliaMiner. However, all ciliopathy clinical features are available on the “Symptoms and Diseases” page. These symptoms are assigned to organ symbols and presented on the same disease symptom summary panels on the “Ciliopathy Names” page.

### Classification of diseases

All disease searches combined with symptoms and subcellular localization data yield a list of 55 different diseases and over 4000 clinical symptoms. The ciliopathy disorders exhibit a variety of diverse symptoms that vary between ciliopathies as well as among the same ciliopathies (**Table 1**). For example, many clinical features, including finger abnormalities like polydactyly, brachydactyly, and syndactyly, situs inversus, cerebral and skeletal anomalies are manifested in the most common ciliopathies, including Bardet-Biedl Syndrome (BBS) (13), Meckel-Gruber Syndrome (MKS) (14), Nephronophthisis (NPHP) (15), and Joubert Syndrome (JBTS) (16) while there are disease-specific clinical features, such as the molar tooth sign for JBTS. We gathered symptoms from all these ciliopathies, collectively labelled as “core ciliopathies”, as a starting point for developing a list of ciliopathy symptoms. As a result, we categorized the most prevalent ciliopathy symptoms as “core ciliopathy symptoms”. It is worth noting that the core ciliopathy symptoms are only obtained from the primary cilia-related ciliopathies; however, the list of motile ciliopathies and motile ciliopathy-associated symptoms are gathered and listed in a separate subgroup on the web page called “Motile Ciliopathy”.

**Table 1:**
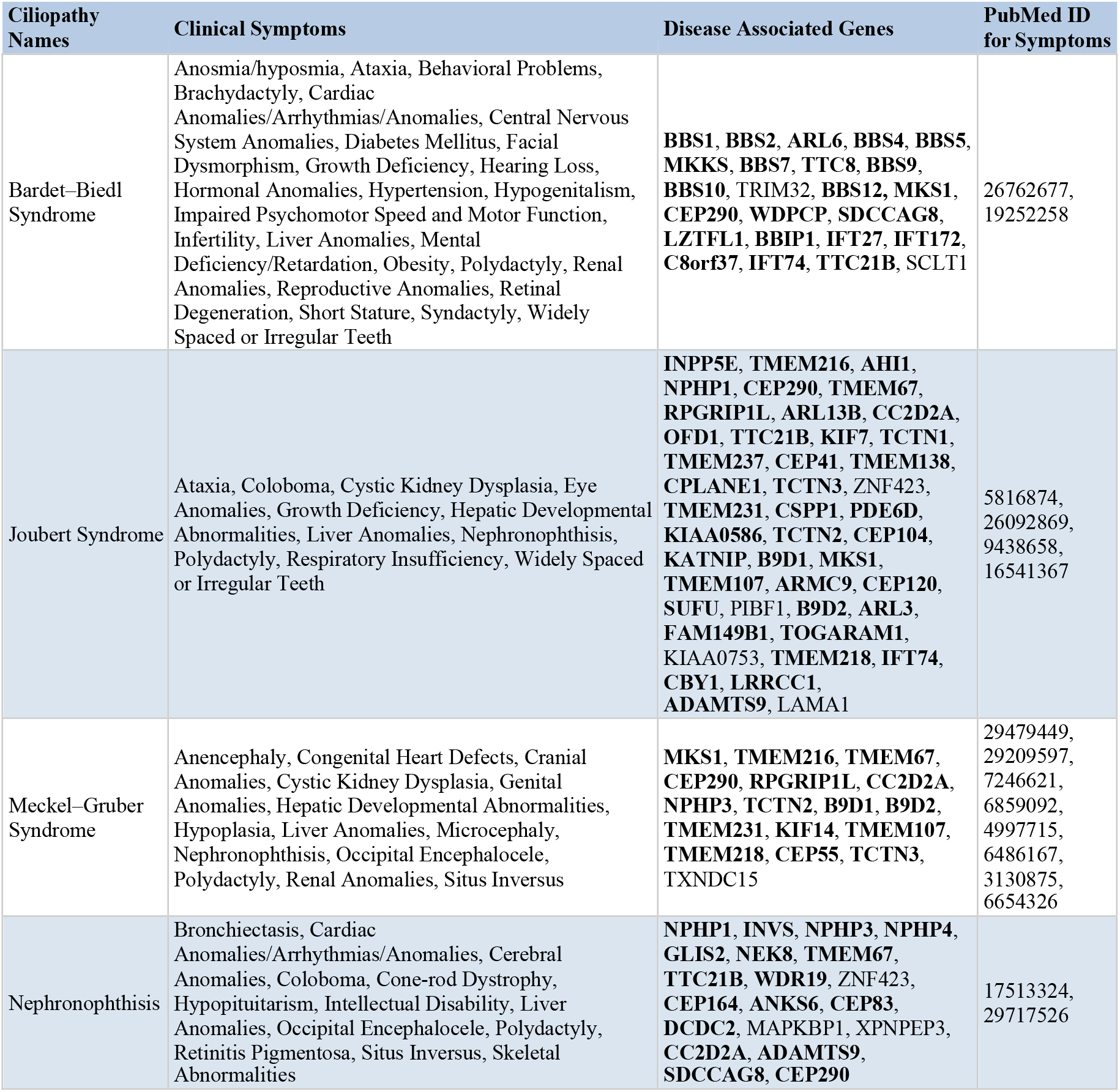
The clinical features and disease-associated genes of selected ciliopathies (non-motile ciliopathies) are shown in the table. The genes in bold are those that have been reported to localize to cilia, while the other genes are not.

Reiter et al. offered the following forms of disease classification for ciliopathies: first-order and second-order ciliopathies (17). Second-order ciliopathies are caused by genes encoding non-cilia localizing proteins, but displaying cilia-related functions, including building cilia or regulating cilia function. First-order ciliopathies are diseases caused by genes encoding proteins localizing to cilia and cilia-related compartments. Even though this review offers an in-depth analysis of both motile and non-motile ciliopathies, the clinical characteristics of disorders were not taken into consideration. However, the disease classification should also consider the clinical features of diseases, because even though proteins may localize to the cilia or ciliary compartments, various clinical symptoms unrelated to ciliopathy may develop as a result of these genes’ non-ciliary functions. For example, Cornelia de Lange syndrome (CdLS), has genes encoding cilia localizing proteins (SMC1A and SMC3) or cilia functioning proteins (HDAC8) (25, 26), CdLS is not yet recognized as a ciliopathy. Because both SMC3 and SMC1A are members of the cohesin complex that holds sister chromatids together, it is currently unknown what symptoms in CdLS are caused by *SMC1A* and *SMC3* functions in cilia (28). It is noteworthy that disease classifications should not focus solely on the core ciliopathy symptoms, because this will lead to mistakes in disease classification. Polycystic kidney disease (PKD), for example, does not have many core ciliopathy symptoms, but it is a well-known ciliopathy. Furthermore, some ciliopathies, such as retinal ciliopathies and renal ciliopathies, cause organ specific symptoms.

For users to compare, the CiliaMiner provides clinical features, subcellular protein localization, and disease-specific clinical features, and uses heatmaps and interactive tables to display all of this data. Similar to Reiter et al., we considered subcellular localization and ciliary functions for disease classification, but clinical characteristics were also taken into account in disease classification. Based on our classification, the diseases are divided into primary, secondary, and atypical ciliopathies. The following criteria were used to determine whether a disease is a primary ciliopathy: 1) comparing disease clinical symptoms to core ciliopathy symptoms; and 2) the localization of disease-causing gene products in cilia and cilia-related compartments (basal body, transition zone, and centrosome). Furthermore, if a disease had previously been proposed as a ciliopathy, we included it in the primary ciliopathy disease without considering further evidence. Secondary diseases are documented by independently checking clinical features and subcellular localization data of condition-associated genes. The disease is listed as a secondary disease if it presents similarity with core clinical symptoms but lacks a protein localizing to the cilia and cilia-related compartments, and vice versa. Finally, many single gene disorders have not been classified into any BBS; NPHP, or other types of ciliopathies but have been reported to be atypical ciliopathies, and these atypical ciliopathies are collected directly from research articles.

Primary ciliopathy consists of 34 diseases (**Supplementary Table 1**), while secondary disease includes 18 diseases (**Supplementary Table 2**). Whole diseases are listed in the tables with disease-associated genes and some clinical features. Bold lettering is used in tables to indicate ciliary proteins which are localized in the cilia and cilia-related compartments as well as all clinical symptoms that are common to all diseases when compared to core ciliopathies.

Atypical ciliopathies were created using the “ciliopathy” keyword search in PubMed between 2018-2022. CiliaMiner displays unclassified ciliopathy genes along with their ciliopathy groups. The atypical ciliopathy genes were reported as ciliopathy genes, and the disease gene names and related reference papers are provided on CiliaMiner. Furthermore, to provide candidate ciliary genes for ciliopathy diseases, we downloaded the CiliaCarta (18) gene list and listed their ciliary localization or cilia-related function papers on CiliaMiner. It is worth noting that our “ciliopathy” or “cilia” search yielded a list of ciliary genes, which were included in the potential ciliopathy gene list on CiliaMiner.

### Database implementation

CiliaMiner is a novel, user-friendly database that provides an up-to-date list of ciliopathy diseases and disease genes, as well as detailed clinical symptoms and other disease-related information. R Shiny (v1.7.1) (19) was used to generate the CiliaMiner website. In addition to the Shiny package, other main libraries like DT (v0.20) (20), ggplot2 (v3.3.5) (21), heatmaply (v1.3.0) (22), and plotly (v4.10.0) (23) are used for generating visual representations. The DT package is employed for ordering, searching, and creating data tables in the user interface. ggplot2 is used for creating graphical representations of the number of ciliopathy genes and their localization presentation on the home page. Plotly and heatmaply are used for creating interactive figures comprising interactive heatmaps and data tables.

Our GitHub repository contains a complete list of the manually curated data along with detailed excel files for each sub-panel (https://github.com/thekaplanlab/CiliaMiner).

## Results

### Database overview and statistics

The last two decades have seen a dramatic expansion of the number of ciliopathies and ciliopathies-associated genes. Even though all relevant data can be found in multiple databases and supplementary files, there is an urgent need for a single and comprehensive database for ciliopathies and ciliopathies-associated genes, so users can access the following: 1. A detailed list of ciliopathies and potential ciliopathies, including symptoms and gene names 2. The updated list of the ciliopathy genes and ciliopathy candidate genes. We, therefore, introduce a new ciliopathy database, called CiliaMiner, which presents a total of 507 genes for different types of ciliopathies based on our disease classification approach. Users may access the CiliaMiner at https://kaplanlab.shinyapps.io/ciliaminer.

Primary ciliopathy has a total of 230 genes associated with 34 different ciliopathy diseases. To assist users, the database includes disease names (also alternative disease names), the names of disease-associated genes as well as the symptoms of each condition. Furthermore, the subcellular localization of proteins encoded by disease-causing genes is supplied for each gene, along with appropriate references, allowing users to view where the products of disease-causing genes are located in cells. The data obtained demonstrated that the protein products of 72, 52, and 28 ciliopathy-related genes, respectively, are found in cilia, the basal body, and the transition zone. 78 ciliopathy encoding proteins do not seem to localize to the cilia-related compartments, but the subcellular localization of data for these genes are collected from the ProteinAtlas, and these proteins localize in the different subcellular departments. Some of the proteins are localized in the plasma membrane (CC2D1A), mitochondria (PAM16, GPX4, XPNPEP3), endoplasmic reticulum (GANAB, DNAJB11, ALG9, LRAT), Golgi apparatus (LRAT, TXNDC15, PI4KB), and centrosome (KIAA0753, LRRCC1) **(Figure 2A)** (24). Additionally, some diseases do not have published genes; just, only the disease condition is known.

**Figure 2:**
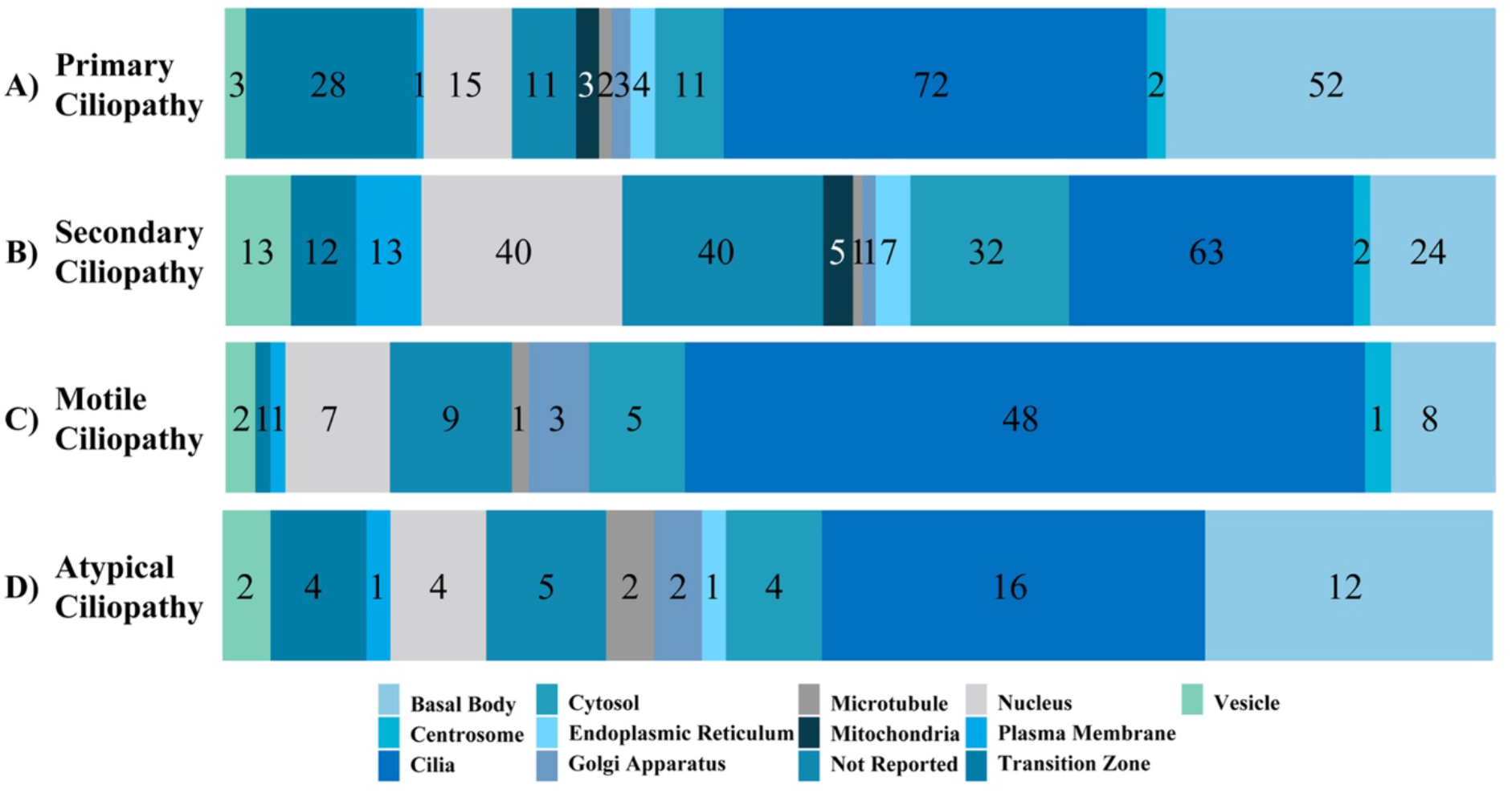
The subcellular localization of ciliopathy disease genes within cells is displayed. Numbers in the bar graph demonstrate how the proteins encoded by the primary, secondary, and atypical ciliopathies-causing genes are localized throughout the cell.

The secondary disease group has 202 genes belonging to 18 secondary ciliopathy diseases. The subcellular localization of proteins encoded by disease-causing genes revealed that 12 proteins are found in the transition zone, 24 in the basal body, and 63 in the cilia; the localization of the rest of the secondary disease-associated genes are localized in the other subcellular departments like plasma membrane (PITPNM3, GUCA1A, MERTK, KLHL7, RGR, CNGA1, DHDDS, SLC7A14, GPC3, SPTBN2, eEF2, TDP1, CC2D1A), mitochondria (IDH3B, HK1, IDH3A, AFG3L2, COA7), endoplasmic reticulum (PNPLA6, CLCC1, MERTK, REEP6, CACNA1A, ELOVL5, PLD3), Golgi apparatus (DRAM2) **(Figure 2B)** (24).

Motile ciliopathies have three diseases: primary ciliary dyskinesia, Birt-Hogg-Dubé syndrome, and Juvenile myoclonic epilepsy. This group presents 75 established disease-associated genes for all. Forty-eight are localized in the cilia, eight are localized in the basal body, and one is found in the transition zone. Other genes are located in the out of the cilia like plasma membrane (GABRA1), Golgi apparatus (GOLGA3, TP73, GABRD), cytosol (DNAAF2, SPAG1, CFAP54, DNAH7, CLCN2). The number of all genes is given in **Figure 2C**.

In addition to primary and secondary diseases, all atypical and potential ciliopathy lists are gathered and presented in the databases. There are 42 genes in the “Atypical Ciliopathy” sub-tab, of which we present unclassified ciliopathy-associated genes **(Figure 2D)**. 274 ciliary proteins encoding genes were carefully gathered, and a list of these hitherto unconnected to ciliopathy genes, as well as data on subcellular localization, is presented in CiliaMiner. We consider each of them as a possible ciliopathy candidate gene. In addition, the list of CilaCarta 934 genes has been included in CiliaMiner for users to examine under the “Potential Ciliary Genes” sub-tab. It is noteworthy that of these genes, 505 of them localize to cilia and cilia-related compartments. Comparing the genes that cause primary, secondary, and atypical ciliopathies demonstrates that numerous genes are responsible for multiple different diseases. A Venn diagram of the common and group-specific genes of primary, secondary, and atypical ciliopathies is shown in **Figure 3**. All ciliopathy subgroups share the gene TTC21B, and both primary and secondary ciliopathies share the genes IFT140, BBS1, BBS2, ARL6, TTC8, CEP290, IFT172, C8orf37, IFT43, GLI3, INPP5E, OFD1, SUFU, ARL3, CC2D1A, GUCY2D, RPE65, RPGRIP1, CRX, CRB1, IMPDH1, RDH12, TULP1, PRPH2, CEP55, CEP83, DCDC2, and DYNC2I2. CFAP45 and PROM1 are found in both atypical and secondary ciliopathies, whereas TTC26, SCLT1, DNAJB11, DYNC2LI1, ALMS1, IFT80, and IFT74 are shared between primary and atypical ciliopathies **(Figure 3)**.

**Figure 3:**
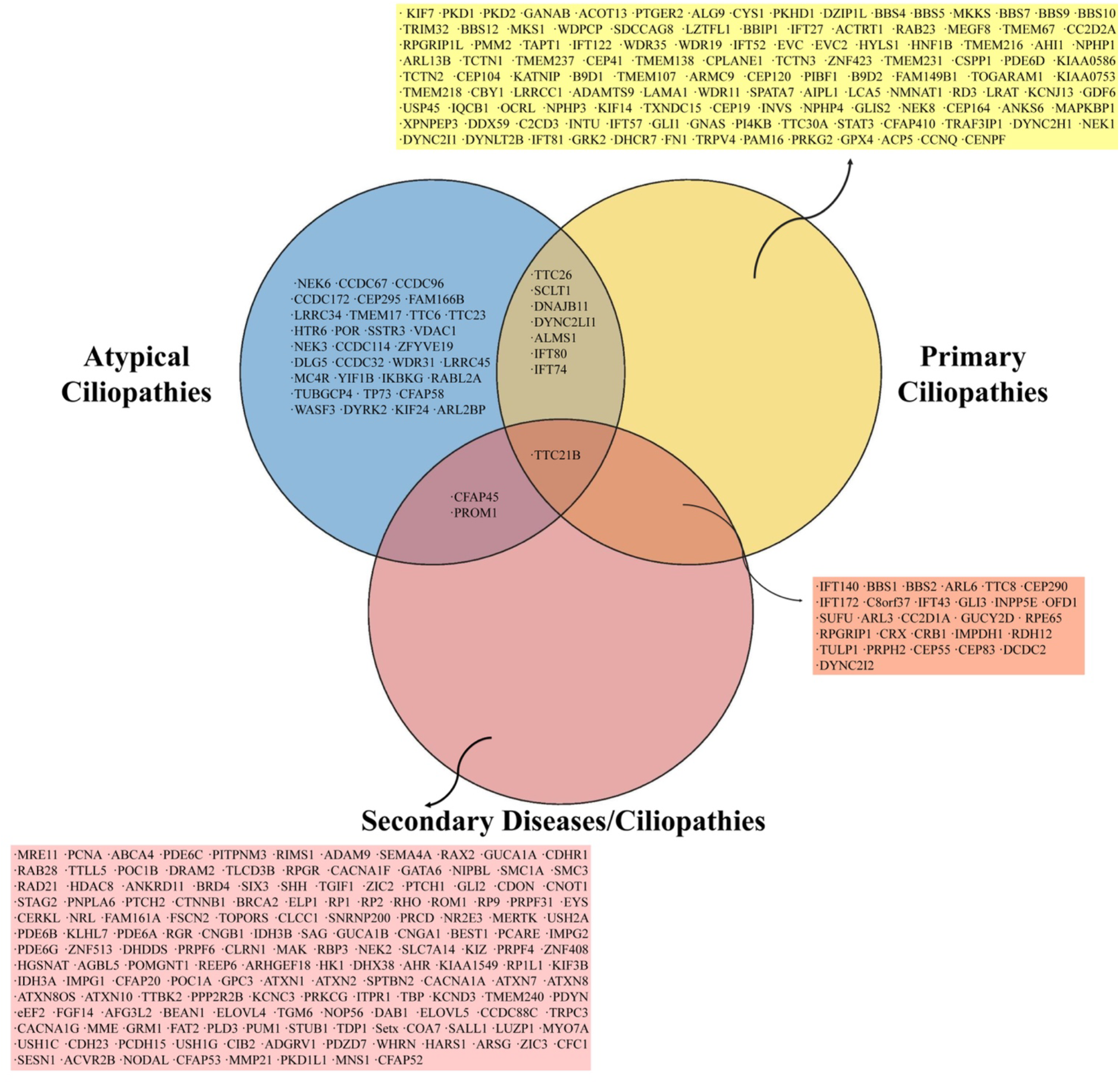
The Venn diagram shows shared genes among ciliopathy-related genes of primary, secondary, and atypical ciliopathies.

For primary ciliopathy, our collection reveals a total of 2354 distinct clinical features, compared to 1784 for secondary ciliopathy. Additionally, motile ciliopathy presents 341 clinical symptoms. We feel that a supplementary table will be outdated if we provide one after some time, so we direct users to the CiliaMiner website for the regularly updated list. All lists can be downloaded from the website. The website was created in a way that users may either search for a symptom name to list diseases where it is present or list all symptoms of a condition. Even for the same condition, distinct symptoms have been recorded for different genes; as a result, the database includes a screening of gene-specific symptoms, enabling users to look for gene-specific symptoms.

### User interface and reactivity

The CiliaMiner homepage provides two search options. Users can do searches using either a disease name or a gene name. Gene Search queries can include human gene names, gene IDs, and Ensembl IDs. On the menu item “Symptoms and Diseases,” there is also an option to search by symptom names. Human gene names, gene IDs, and Ensembl IDs can all be used in Gene Search queries. Gene names, gene IDs, OMIM numbers, ciliopathy names, and localization references can all be utilized to do specialized searches across all pages and sub-tabs.

CiliaMiner has different visualization tools for the comparative analysis of ciliopathies and clinical symptoms. Relative heat maps have been integrated into the Ciliopathy Names page. Specifically, primary ciliopathies and secondary ciliopathies tabs can compare different ciliopathies regarding clinical symptoms. In addition, heatmaps can be regenerated by user-selected ciliopathies and a graphical representation of user inputs that can be used for comparing cilia-related clinical features.

The representative figures of symptoms are also another visualization way for summarizing ciliopathies. This visualization of symptoms was created by using symptom supergroups to understand ciliopathies’ effect on organs and systems in the human body. These 16 supergroups; aural, neural, ophthalmic, skeletal, respiratory, hormonal, reproductive, facial, cerebral, renal, coronary and vascular, nasal, liver, cognitive, digestive, and organ anomalies were created for a straightforward understandable clinical representation by using ciliopathy based clinical symptoms.

### Strengths of CiliaMiner

The continuing increases in the number of ciliopathies and ciliopathy genes present enormous challenges for clinical and basic scientists. They need to visit multiple databases and look up specific information. In this regard, CiliaMiner, a novel manually-curated database for ciliopathy, will be helpful since it makes all ciliopathies, ciliopathy genes, disease- and gene-specific symptoms, and prospective ciliopathy candidate genes easily accessible. Furthermore, while providing a thorough list of well-known ciliopathies, CiliaMiner also lists a potential ciliopathy candidate, such as Cornelia de Lange syndrome. Cornelia de Lange syndrome (CdLS) has seven disease-associated genes; 2 of them localize (*SMC1A* and *SMC3*) to cilia (25), and 2 of them (*ANKRD11* and *HDAC8*) are implicated in cilia (26, 27). Although the precise relationship between CdLS and cilia is not yet understood, localization and functional data suggest that several symptoms, including hearing loss, abnormal hands and limbs, and cardiac issues, may be brought on by the cilia-related functions of these genes. Additionally, the network analysis indicates CdLS disease genes interact with many ciliary genes **(Figure 4)**. To establish a connection between cilia and these genes, more work is required.

**Figure 4:**
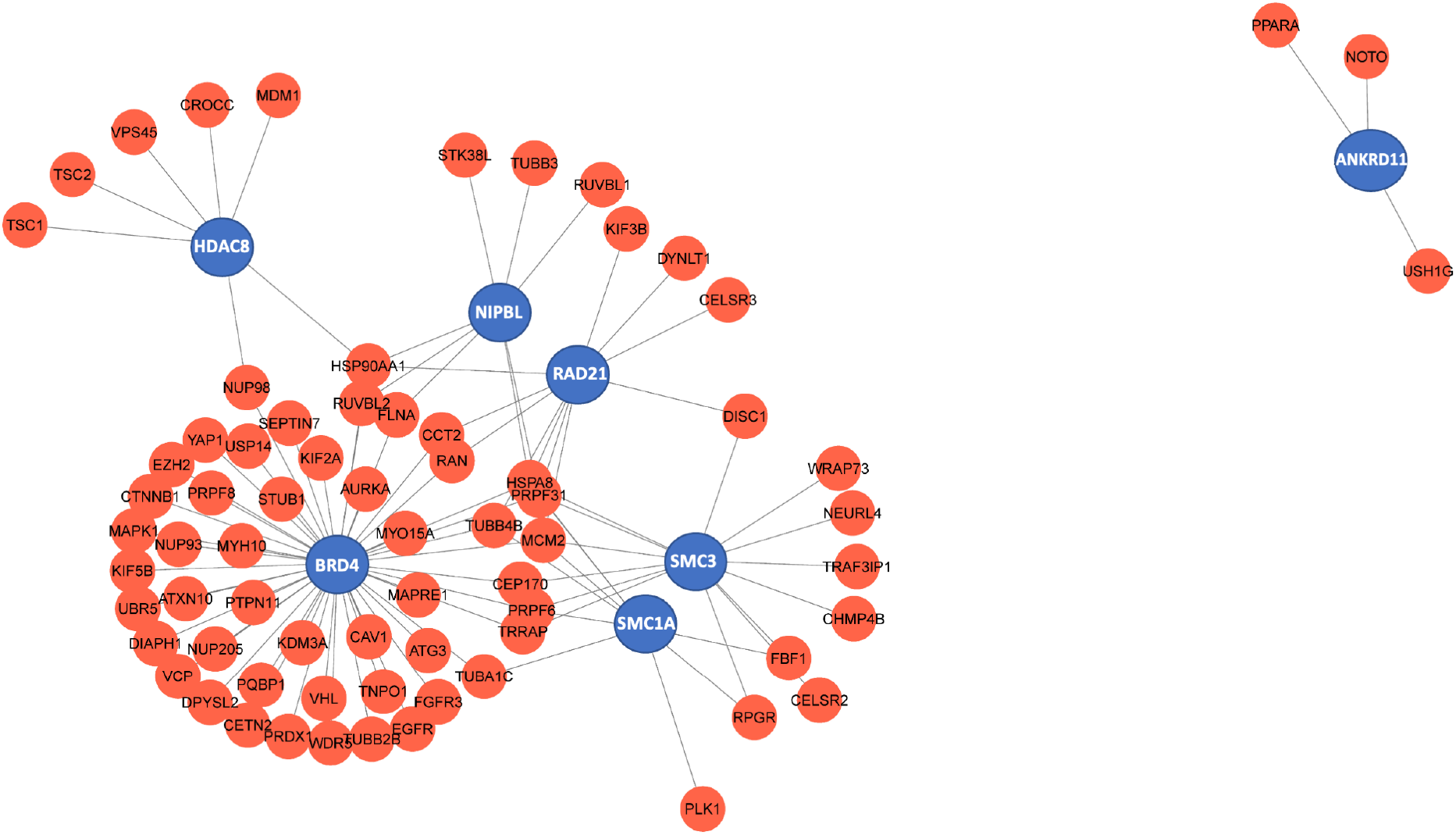
Genes that cause Cornelia de Lange syndrome (CdLS) were analysed using a network. Red represents the ciliary genes while blue points CdLS associated genes.

Researchers that work with model species are crucial to the cilia field, and CiliaMiners makes it simple to find the orthologs of genes that cause disorders. The database enables searches for mouse, zebrafish, clawed frog, fruit fly, and worm orthologs of human genes linked to diseases. In conclusion, CiliaMiner is an comprehensive database for the exploration of cilia fields, offering a detailed list of ciliopathies, ciliopathy genes, clinical characteristics of each illness, and prospective ciliopathy candidate genes. The content will be constantly updated and users will be able to add and/or correct the relevant information on the website.

---

**Supplementary Table 1**: The clinical features and published genes of all primary ciliopathies. Only the genes shown in bold localize to cilia; all other genes do not. Clinical characteristics related with ciliopathies are shown in bold.

**Supplementary Table 2**: The clinical features and established genes of all secondary diseases. The only genes found to localize to cilia are those in bold. Clinical features associated with ciliopathies are highlighted (bold).

## Acknowledgments

We thank Associate Professor Duygu Sacar for allowing Mehmet Emin Orhan to help this project. We appreciate Mustafa Pir’s assistance with Figure 4. His coding and CilioGenics work enable us to produce Figure 4. We thank Dr. Abdullah Sezer for critical reading of the manuscript.

