## Supplementary Table 1 for "CiliaMiner: an integrated database for Ciliopathy Genes and Ciliopathies"

**Table 2:** The clinical features and published genes of all primary ciliopathies.

| **Ciliopathy Names** | **Clinical Symptoms** | **Ciliopathy Associated Genes** | **PubMed ID for Symptoms** |
| --- | --- | --- | --- |
| Acrocallosal Syndrome | Agenesis or Partial Agenesis of Corpus Callosum, Brachydactyly, **Cranial Anomalies**, Frontal Bossing, Hypertelorism, **Impaired Psychomotor Speed and Motor Function**, Macrocephaly, **Mental Deficiency/Retardation**, **Polydactyly** | **KIF7** | 31399769, 16582532,  2658584 |
| Al-Gazali-Bakalinova Syndrome | Flat Malar Regions, Frontal Bossing,  Hypertelorism, Low-set Ears, Macrocephaly, Multiple Epiphyseal Dysplasia, **Short Stature** | **KIF7** | 22587682, 23431744 |
| Alström Syndrome | **Cardiac**  **Anomalies/Arrhythmias/Anomalies, Cone-rod Dystrophy**, **Diabetes Mellitus**, **Hearing Loss**, **Hypogenitalism**, Insulin Resistance, **Liver Anomalies**, Nystagmus, **Obesity**, Photophobia/Photosensitivity, Polycystic Ovary Syndrome, **Renal Anomalies**, **Short Stature** | **ALMS1** | 30578508 |
| Autosomal Dominant Polycystic Kidney Disease | Aneurysm, **Cerebral Anomalies**, Colonic Diverticula, **Cystic Kidney Dysplasia, Hypertension**, Polycystic Liver, **Renal Anomalies** | **PKD1, PKD2,**  GANAB**,** DNAJB11**, IFT140,** ACOT13**,** PTGER2**,** ALG9**, CYS1** | 13723091, 1670785,  6766288, 6500563,  1670785, 688689,  1583643 |
| Autosomal Recessive Polycystic Kidney Disease | Aneurysm, **Cerebral Anomalies**, Colonic  Diverticula, **Cystic Kidney Dysplasia, Hypertension**, Polycystic Liver, **Renal Anomalies** | **PKHD1, DZIP1L** | 13723091, 1670785,  6766288, 6500563,  1670785, 688689,  1583643 |
| Bardet–Biedl Syndrome | Anosmia/hyposmia, **Ataxia**, **Behavioral**  **Problems**, **Brachydactyly**, **Cardiac Anomalies/Arrhythmias/Anomalies**, **Central Nervous System Anomalies**, **Diabetes Mellitus**, **Facial Dysmorphism, Growth Deficiency**, **Hearing Loss**, **Hormonal Abnomalies**, **Hypertension**, **Hypogenitalism**, **Impaired Psychomotor Speed and Motor Function**, **Infertility**, **Liver Anomalies**, **Mental Deficiency/Retardation**, **Obesity**, **Polydactyly**, **Renal Anomalies**, Reproductive Anomalies, **Retinal Degeneration**, **Short Stature**, **Syndactyly**, **Widely Spaced or Irregular Teeth** | **BBS1, BBS2, ARL6,**  **BBS4, BBS5, MKKS, BBS7, TTC8, BBS9, BBS10,** TRIM32**, BBS12, MKS1, CEP290, WDPCP, SDCCAG8, LZTFL1, BBIP1, IFT27, IFT172, C8orf37, IFT74, TTC21B,** SCLT1 | 26762677, 19252258 |
| Bazex-Dupré-Christol Syndrome | **Cardiac**  **Anomalies/Arrhythmias/Anomalies**, **Cerebral Anomalies, Diabetes Mellitus**, **Eye Anomalies**, **Facial Dysmorphism, Genital Anomalies**, **Hormonal Abnomalies**, Hypotrichosis, **Liver Anomalies,** Neurological Problems, **Polydactyly**, **Situs Inversus,** Sparse Hair, Unbalanced Pigmentation, Visceral Malignancies | **ACTRT1** | 7747764, 708616,  8456866, 16120174,  1642265, 29808590 |
| Bilateral Polycystic Kidney  Disease | Aneurysm, **Cerebral Anomalies**, Colonic  Diverticula, **Cystic Kidney Dysplasia,** | **TTC21B** | 13723091, 1670785,  6766288, 6500563, |

|  | **Hypertension**, Polycystic Liver**, Renal Anomalies** |  | 1670785, 688689,  1583643 |
| --- | --- | --- | --- |
| Biliary, Renal, Neurologic, and Skeletal Syndrome | **Cardiac**  **Anomalies/Arrhythmias/Anomalies**, **Cerebral Anomalies**, Bile Duct System Anomalies, **Diabetes Mellitus**, **Eye Anomalies, Facial Dysmorphism**, Hydrocephalus, **Liver Anomalies**, **Polydactyly**, **Situs Inversus** | **TTC26** | 31595528, 24596149,  34177428 |
| Caroli Disease | Bile Duct System Anomalies, **Liver**  **Anomalies** | **PKHD1** | 17461492, 11337358 |
| Carpenter Syndrome | Acrocephaly, **Brachydactyly**, **Cerebral Anomalies**, **Cranial Anomalies**, **Facial Dysmorphism**, **Hypogenitalism**, Limb Anomalies, **Mental Deficiency/Retardation**, **Obesity**,  Polyhydramnios, **Syndactyly** | **RAB23**, MEGF8 | 25168863, 19974019,  5935752 |
| COACH Syndrome | **Ataxia**, **Cerebral Anomalies, Coloboma**,  **Growth Deficiency** | **TMEM67, CC2D2A,**  **RPGRIP1L,** PMM2 | 32032630, 12385458 |
| Complex Lethal Osteochondrodysplasia | **Cerebral Anomalies**, **Cystic Kidney Dysplasia**, **Diabetes Mellitus**, **Growth Deficiency**, **Infertility**, **Obesity**, **Retinal Degeneration**, **Skeletal Abnormalities** | **TAPT1** | 26365339 |
| Cranioectodermal Dysplasia | Anteverted Nares, Arched Palate, **Brachydactyly**, **Congenital Heart Defects,** Everted Lower Lip, Frontal Bossing, **Growth Deficiency**, Hypodontia, Hypoplastic and/or Dysplastic Nails, Hypotelorism, Limb Anomalies, Low-set Ears, **Polydactyly**, **Skeletal Abnormalities,** Small Thorax, Sparse Hair, **Syndactyly**, Telecanthus, **Widely Spaced or Irregular Teeth**,  Widened Nasal Bridge | **IFT122, WDR35,**  **IFT43, WDR19, IFT52, IFT140** | 24027799, 2661822 |
| Ellis-van Creveld Syndrome | Brachydactyly, **Cardiac Anomalies/Arrhythmias/Anomalie**s, Chest Deformity, **Congenital Heart Defects**, **Growth Deficiency**, **Hypertension**, Hypoplastic and/or Dysplastic Nails, **Polydactyly**, **Respiratory Insufficiency**, Short Ribs, Skeletal Dysplasia, **Widely Spaced or Irregular Teeth** | **EVC, EVC2** | 17547743, 27325544,  31850774 |
| Greig Cephalopolysyndactyly Syndrome | Agenesis or Partial Agenesis of Corpus Callosum, **Central Nervous System Anomalies**, Cognitive Impairment, **Eye Anomalies**, Frontal Bossing, **Growth Deficiency**, Hernia, Hypertelorism, Macrocephaly, **Polydactyly**, Seizures,  **Syndactyly**, Widened Nasal Bridge | **GLI3** | 20301619, 12794692,  12794692, 18435847 |
| Hydrolethalus Syndrome | Anomalies of The Lower Jaw, **Central Nervous System Anomalies**, **Congenital Heart Defects**, Holoprosencephaly, Hydrocephalus, Limb Anomalies, Organ Dysfunction, **Polydactyly**, **Respiratory**  **Insufficiency** | **HYLS1, KIF7** | 7028327, 2407847,  11152149 |
| Infantile Polycystic Kidney Disease | Aneurysm, **Cerebral Anomalies**, Colonic Diverticula, **Cystic Kidney Dysplasia, Hypertension**, Polycystic Liver, **Renal**  **Anomalies** | HNF1B | 13723091, 1670785,  6766288, 6500563,  1670785, 688689,  1583643 |

| Joubert Syndrome | **Ataxia**, **Coloboma**, **Cystic Kidney Dysplasia**, **Eye Anomalies**, **Growth Deficiency**, **Hepatic Developmental Abnormalities, Liver Anomalies**, **Nephronophthisis**, **Polydactyly**, **Respiratory Insufficiency, Widely Spaced or Irregular Teeth** | **INPP5E, TMEM216,**  **AHI1, NPHP1, CEP290, TMEM67, RPGRIP1L, ARL13B, CC2D2A, OFD1, TTC21B, KIF7, TCTN1, TMEM237, CEP41, TMEM138, CPLANE1, TCTN3,** ZNF423**, TMEM231, CSPP1, PDE6D, KIAA0586, TCTN2, CEP104, KATNIP, B9D1, MKS1, TMEM107, ARMC9, CEP120, SUFU,** PIBF1**, B9D2, ARL3, FAM149B1, TOGARAM1,** KIAA0753**, TMEM218, IFT74, CBY1,** CC2D1A**,** LRRCC1**, ADAMTS9,** LAMA1 | 5816874, 26092869,  9438658, 16541367 |
| --- | --- | --- | --- |
| Kallmann Syndrome | Anophthalmia, Delayed Puberty,  Holoprosencephaly, Hypotelorism, **Infertility**, **Microcephaly**, Microphthalmia, **Obesity** | **WDR11** | 29263200 |
| Leber Congenital Amaurosis | Absence of The Pupillary Reflex, **Eye Anomalies**, Eye Rubbing or Poking, **Intellectual Disability**, Nystagmus, Photoreceptors Dysfunction, Visual Loss | **GUCY2D**, RPE65,  **SPATA7**, **AIPL1**, **LCA5**, **RPGRIP1**, CRX, CRB1, **NMNAT1**, **CEP290**, IMPDH1, RD3, RDH12, LRAT, **TULP1**, KCNJ13, GDF6, **PRPH2**, USP45, **IQCB1**, **INPP5E** | **31639339, 30578499,**  **31009524** |
| Lowe Oculocerebrorenal Syndrome | **Behavioral Problems**, **Eye Anomalies,**  **Growth Deficiency**, **Intellectual Disability**, Urinary Anomalies | **OCRL** | 27011217, 20301653 |
| McKusick-Kaufman Syndrome | **Cardiac**  **Anomalies/Arrhythmias/Anomalies**, Hydrometrocolpos, **Polydactyly** | **MKKS** | 14172277, 21954533,  8209897 |
| Meckel–Gruber Syndrome | Anencephaly, **Congenital Heart Defects**, **Cranial Anomalies**, **Cystic Kidney Dysplasia**, **Genital Anomalies, Hepatic Developmental Abnormalities**, **Hypoplasia**, **Liver Anomalies**, **Microcephaly**, **Nephronophthisis**, **Occipital Encephalocele**, **Polydactyly**, **Renal Anomalies**, **Situs Inversus** | **MKS1, TMEM216,**  **TMEM67, CEP290, RPGRIP1L, CC2D2A, NPHP3, TCTN2, B9D1, B9D2, TMEM231, KIF14, TMEM107, TMEM218, CEP55, TCTN3,** TXNDC15 | 29479449, 29209597,  7246621, 6859092,  4997715, 6486167,  3130875, 6654326 |
| Morbid Obesity and Spermatogenic Failure | **Hypertension**, Insulin Resistance, **Liver**  **Anomalies**, **Obesity**, Spermatogenic Failure | **CEP19** | 24268657 |
| Nephronophthisis | **Bronchiectasis**, **Cardiac**  **Anomalies/Arrhythmias/Anomalies**, **Cerebral Anomalies**, **Coloboma**, | **NPHP1, INVS,**  **NPHP3, NPHP4, GLIS2, NEK8,** | 17513324, 29717526 |

|  | **Cone-rod Dystrophy**, **Hypopituitarism**, **Intellectual Disability**, **Liver Anomalies, Occipital Encephalocele**, **Polydactyly**, **Retinitis Pigmentosa**, **Situs Inversus, Skeletal Abnormalities** | **TMEM67, TTC21B,**  **WDR19,** ZNF423**, CEP164, ANKS6, CEP83, DCDC2,** MAPKBP1**,** XPNPEP3**, CC2D2A, ADAMTS9, SDCCAG8, CEP290** |  |
| --- | --- | --- | --- |
| Orofaciodigital Syndrome | Brachydactyly, **Cardiac**  **Anomalies/Arrhythmias/Anomalies**, **Eye Anomalies**, Hydrocephalus, Hypertelorism, Low-set Ears, **Polydactyly**, **Skeletal Abnormalities**, Tongue Anomalies | **OFD1, TCTN3,**  DDX59**, CPLANE1, C2CD3,** KIAA0753**, TMEM107, INTU, IFT57** | 19396822, 28289185 |
| Polycystic Kidney Disease | Aneurysm, **Cerebral Anomalies**, Colonic  Diverticula, **Cystic Kidney Dysplasia**, **Hypertension**, Polycystic Liver, **Renal Anomalies** | **GLI1,** PMM2**, GNAS,**  PI4KB**, TTC30A,**  STAT3 | 13723091, 1670785,  6766288, 6500563,  1670785, 688689,  1583643 |
| Renal-hepatic-pancreatic Dysplasia | **Cerebral Anomalies**, Dandy–Walker  Malformation, Gastrointestinal Dysfunction, **Hypoplasia**, **Liver Anomalies,** Pancreatic Fibrosis, **Renal Anomalies**, Situs Abnormalities | **NPHP3, NEK8,**  DNAJB11 | 17605805, 32341812 |
| Retinal Dystrophy | Brachydactyly, **Cone-rod Dystrophy,**  Night Blindness, **Obesity**, Retinal Dystrophy, **Short Stature**, Small Thorax, Visual Loss | **CFAP410, CEP83,**  **TTC21B** | 27548899, 26294103 |
| RHYNS Syndrome | Growth Hormone Deficiency,  **Hypopituitarism**, **Nephronophthisis**, **Retinitis Pigmentosa**, Skeletal Dysplasia, Upper Eyelid Ptosis | **TMEM67** | 11391657, 29891882,  9375913 |
| Senior-Løken Syndrome | Chronic Tubulointerstitial Nephritis,  **Cystic Kidney Dysplasia**, **Hypertension**, **Nephronophthisis**, Nystagmus, Photophobia/Photosensitivity, Reduced Concentrating Ability | **NPHP1, NPHP4,**  **IQCB1, CEP290, SDCCAG8, WDR19, TRAF3IP1, NPHP3** | 30578507, 22819833 |
| Short-rib Thoracic Dysplasia | Brachydactyly, Gastrointestinal Dysfunction, **Hypoplasia**, Limb Anomalies, Organ Dysfunction, **Renal Anomalies, Respiratory Insufficiency**, **Retinal Degeneration** | **IFT80, DYNC2H1,**  **TTC21B, WDR19, NEK1, WDR35, DYNC2I1, IFT140, IFT172, DYNC2I2, CEP120, KIAA0586, DYNC2LI1, IFT52, DYNLT2B, IFT43, IFT81, INTU,** KIAA0753**,** GRK2 | 23985472, 23456818 |
| Smith-Lemli-Opitz Syndrome | **Cardiac**  **Anomalies/Arrhythmias/Anomalies, Facial Dysmorphism**, **Mental Deficiency/Retardation**, Micrognathia, Photophobia/Photosensitivity, **Syndactyly**, Upper Eyelid Ptosis | **DHCR7** | 9714007, 9714006,  9678700 |
| Spondylometaphyseal Dysplasia | Chest Deformity, **Eye Anomalies**,  **Hypoplasia**, Platyspondyly, **Retinitis Pigmentosa**, Short Ribs, **Short Stature,** Skeletal Dysplasia, Small Thorax, Visual Loss | **CFAP410,** FN1**,**  **TRPV4,** PAM16**,** PRKG2**,** GPX4**,** ACP5 | 28123176, 9266195,  21910225 |
| STAR Syndrome | Atresia, **Congenital Heart Defects**, **Eye**  **Anomalies**, **Growth Deficiency**, **Renal Anomalies**, **Reproductive Anomalies**, | CCNQ | 28225384 |

|  | **Skeletal Abnormalities**, **Syndactyly**, Telecanthus |  |  |
| --- | --- | --- | --- |
| Stromme Syndrome | **Cranial Anomalies**, **Eye Anomalies**,  Gastrointestinal Dysfunction,  **Microcephaly** | **CENPF** | 28401041, 8261651 |
| Weyers Acrofacial Dysostosis | Anomalies of The Lower Jaw, Hypoplastic and/or Dysplastic Nails, **Polydactyly**, Prominent Ear Antihelices, **Short Stature**, **Widely Spaced or Irregular Teeth** | **EVC, EVC2** | 12989297, 6499270 |
