## Supplementary Table 2 for "CiliaMiner: an integrated database for Ciliopathy Genes and Ciliopathies"

**Table 3:** The clinical features and established genes of all secondary diseases.

| **Ciliopathy Names** | **Clinical Symptoms** | **Disease Associated Genes** | **PubMed ID for Symptoms** |
| --- | --- | --- | --- |
| Ataxia-telangiectasia-like Disorder | **Cerebral Anomalies**, **Eye Anomalies**,  Immunodeficiency, Loss of Balance and Coordination, Sensitivity To Ionising Radiation | MRE11, PCNA | 15279810, 29170652,  27884168 |
| Cone-Rod Dystrophy | Color Vision Deficit, Decreased Central Vision, Hemeralopia, Loss of Peripheral Field, Night Blindness, Photophobia, Reading Difficulties, Visual Loss | CRX, ABCA4, **PDE6C**,  PITPNM3, **GUCY2D**, RIMS1, **ADAM9**, SEMA4A, RAX2, P**ROM1**, **RPGRIP1**, GUCA1A, **CDHR1**, **C8orf37**, **RAB28**, **TTLL5**, **POC1B**, DRAM2, TLCD3B, **RPGR**, CACNA1F, **DYNC2I2** | 30578485, 29555955 |
| Congenital Heart Disease | **Cardiac**  **Anomalies/Arrhythmias/Anomalies**, Fatigue, **Hypertension**, Limited Exercise Ability, **Respiratory Insufficiency** | GATA6 | 25638345, 29400306 |
| Cornelia de Lange Syndrome | **Cardiac**  **Anomalies/Arrhythmias/Anomalies**, **Facial Dysmorphism**, Gastrointestinal Dysfunction, Genitourinary Malformations, **Growth Deficiency**, Hernia, Hirsutism, Hypertrichosis, **Intellectual Disability**, Limb Anomalies, **Microcephaly**, Postnatal Growth Failure/Retardation, Prenatal Growth Retardation, Pyloric Stenosis, **Widely Spaced or Irregular Teeth** | NIPBL, **SMC1A**, **SMC3**, RAD21, HDAC8, ANKRD11, BRD4 | 25209348, 20301283 |
| Holoprosencephaly | Cleft Lip, Cleft Palate, **Cranial**  **Anomalies**, Forebrain Malformation, Hypotelorism, **Microcephaly**, Widened Nasal Bridge | SIX3, **SHH**, TGIF1,  ZIC2, **PTCH1**, **GLI2**, CDON, CNOT1, STAG2 | 33168217, 29770994 |
| Laurence–Moon Syndrome | Growth Hormone Deficiency,  **Hypogenitalism**, **Mental Deficiency/Retardation**, Proximal Placement of The Thumb, **Retinal Degeneration** | PNPLA6 | 8053762 |
| Medulloblastoma | **Ataxia**, **Eye Anomalies**, Hydrocephalus,  Loss of Balance and Coordination, Nausea, Vomiting | **PTCH2**, **CTNNB1**, **SUFU**, BRCA2, ELP1 | 18031705, 8756384 |
| Mental Retardation, Truncal  Obesity, Retinal Dystrophy, and Micropenis | **Genital Anomalies**, **Mental**  **Deficiency/Retardation**, **Obesity**, Retinal Dystrophy | **INPP5E** | 16493448 |
| Multinucleated Neurons,  Anhydramnios, Renal Dysplasia, Cerebellar Hypoplasia, and Hydranencephaly | Absence of The Telencephalon,  Anhydramnios, **Cerebral Anomalies**, **Eye Anomalies**, Reduced Odor Identification, **Renal Anomalies**, Severe Hydranencephaly, **Syndactyly** | **CEP55** | 9359650, 6438176,  3130870 |
| Neonatal Sclerosing Cholangitis | Acholic Stools, Cholestasis,  **Hypertension**, **Liver Anomalies**, Neonatal Icterus, **Renal Anomalies** | **DCDC2** | 27319779 |
| Pallister-Hall Syndrome | Hypothalamic Hamartoma, **Polydactyly** | **GLI3** | 20301638, 21108399 |
| Retinitis Pigmentosa | Blindness, **Central Nervous System**  **Anomalies**, **Growth Deficiency**, Night | **RP1**, **RP2**, **RPGR**,  **RHO**, **ROM1**, **PRPH2**, | 23701314, 25345673 |

|  | Blindness, Photoreceptors Dysfunction, Visual Loss | RP9, IMPDH1, **PRPF31**, CRB1, RDH12, **TULP1**, ABCA4, RPE65, **OFD1**, **EYS**, **CERKL**, NRL, **FAM161A**, **FSCN2**, **TOPORS**, CLCC1, **SNRNP200**, SEMA4A, PRCD, NR2E3, MERTK, **USH2A**, PDE6B, P**ROM1**, KLHL7, PDE6A, RGR, CNGB1, IDH3B, SAG, GUCA1B, CNGA1, BEST1, **TTC8**, **PCARE**, **ARL6**, IMPG2, PDE6G, ZNF513, DHDDS, **PRPF6**, **CLRN1**, **MAK**, **C8orf37**, **CDHR1**, RBP3, **NEK2**, SLC7A14, **KIZ**, **PRPF4**, **IFT172**, ZNF408, HGSNAT, **BBS2**, AGBL5, **POMGNT1**, REEP6, **ARHGEF18**, HK1, **IFT140**, **IFT43**, **ARL3**, DHX38, AHR, **KIAA1549**, **RP1L1**, **KIF3B**, IDH3A, IMPG1, **CEP290**, **CEP83**, **BBS1**, **CFAP20** |  |
| --- | --- | --- | --- |
| Short Stature, Onychodysplasia, Facial Dysmorphism, and Hypotrichosis | Facial Dysmorphism, Frontal Balding,  **Genital Anomalies**, **Growth Deficiency**, Hypertelorism, **Hypoplasia**, Limb Anomalies, Macrocephaly, Macrostomia, **Short Stature**, Small Nose | POC1A | 1632447, 22840364 |
| Simpson-Golabi-Behmel Syndrome | Anomalies of The Lower Jaw, Anteverted  Nares, **Brachydactyly**, **Cardiac Anomalies/Arrhythmias/Anomalies**, Cleft Lip, Overgrowth, **Polydactyly**, Tongue Anomalies, Widened Nasal Bridge | GPC3, **OFD1** | 1227524, 2018065,  6538755, 1456280 |
| Spinocerebellar Ataxia | **Cerebral Anomalies**, Cognitive Impairment, **Hearing Loss**, **Impaired Psychomotor Speed and Motor Function**, Limb Anomalies, Loss of Balance and Coordination, Nystagmus, Ophthalmoplegia, Parkinsonism, Seizures | ATXN1, ATXN2,  SPTBN2, CACNA1A, **ATXN7**, ATXN8, ATXN8OS, **ATXN10**, **TTBK2**, PPP2R2B, **KCNC3**, PRKCG, ITPR1, TBP, KCND3, TMEM240, PDYN, eEF2, FGF14, AFG3L2, BEAN1, ELOVL4, TGM6, NOP56, DAB1, ELOVL5, CCDC88C, TRPC3, CACNA1G, MME, GRM1, FAT2, PLD3, PUM1, STUB1, TDP1, Setx, COA7 | 30284037, 30975995 |

| Townes-Brocks Syndrome | Deformed External Ears, Gastroesophageal Reflux, **Genital Anomalies**, **Hearing Loss**, Hypertelorism, Polyhydramnios, Reading Difficulties, **Reproductive Anomalies**, Visual Loss | SALL1, **LUZP1** | 5042490, 671168,  2667456, 20520617,  8357560, 16088922,  11484202 |
| --- | --- | --- | --- |
| Usher Syndrome | **Cerebral Anomalies**, **Hearing Loss**, **Mental Deficiency/Retardation**, Reduced Odor Identification, **Retinitis Pigmentosa** | **MYO7A**, USH1C,  **CDH23**, **PCDH15**, **USH1G**, **CIB2**, **USH2A**, **ADGRV1**, **PDZD7**, **WHRN**, **CLRN1**, HARS1, ARSG | 16987892 |
| Visceral Heterotaxy | **Congenital Heart Defects**, Situs Ambiguus, **Situs Inversus** | ZIC3, CFC1, SESN1,  ACVR2B, NODAL, **CFAP53**, MMP21, **PKD1L1**, **MNS1**, **CFAP52**, **CFAP45**, **TTC21B**, CC2D1A | 22726404, 3674105,  9354777 |
